# Genome-wide investigation of gene-cancer associations for the prediction of novel therapeutic targets in oncology

**DOI:** 10.1101/2020.01.30.927285

**Authors:** Adrián Bazaga, Dan Leggate, Hendrik Weisser

## Abstract

A major cause of failed drug discovery programs is suboptimal target selection, resulting in the development of drug candidates that are potent inhibitors, but ineffective at treating the disease. In the genomics era, the availability of large biomedical datasets with genome-wide readouts has the potential to transform target selection and validation. In this study we investigate how computational intelligence methods can be applied to predict novel therapeutic targets in oncology. We compared different machine learning classifiers applied to the task of drug target classification for nine different human cancer types. For each cancer type, a set of “known” target genes was obtained and equally-sized sets of “non-targets” were sampled multiple times from the human protein-coding genes. Models were trained on mutation, gene expression (TCGA), and gene essentiality (DepMap) data. In addition, we generated a numerical embedding of the interaction network of protein-coding genes using deep network representation learning and included the results in the modeling. We assessed feature importance using a random forests classifier and performed feature selection based on measuring permutation importance against a null distribution. Our best models achieved good generalization performance based on the AUROC metric. With the best model for each cancer type, we ran predictions on more than 15,000 protein-coding genes to identify potential novel targets. Our results indicate that this approach may be useful to inform early stages of the drug discovery pipeline.

## Introduction

The process of drug discovery is expensive and time-consuming. It is exceedingly difficult to produce an efficacious molecule and gain approval as a drug^1^. Target identification is a pivotal stage early in this process. Having the right target is critical to avoid costly development of a modulator that is in the end ineffective at treating the pathology of interest. Indeed, lack of efficacy has been identified as the main reason for late-stage failure of drug development programs^2^.

A landmark study has highlighted the value of genetic evidence linking a target and disease for drug discovery^3^, estimating that selecting targets supported by genetic data could double the success rate in the clinical development pipeline. Experts routinely consult large biomedical and genomic data resources^4^ to guide target identification and validation. Examples for the field of oncology include The Cancer Genome Atlas (TCGA)^5^ or the Cancer Dependency Map (DepMap)^6^. In their manual analyses, experts typically consider each data source and data type (e.g. mutations and gene expression) independently, and weigh information for each individual source against each other using subjective criteria.

Computational approaches, applicable across all stages of drug discovery and biomedical research, provide cost-effective options to guide experts with data-driven decisions, potentially speeding up the process and reducing failure rates^7^. There is a significant amount of literature concerning the application of machine learning methods in the drug discovery pipeline. These works span application fields such as target identification, target-disease association, drug design, drug repurposing, patient stratification, and biomarker discovery^7,8^. In the novel target identification field, Kumari et al.^9^ proposed an improved random forest (RF) algorithm that integrates bootstrap and rotation feature matrix components, to discriminate human drug targets from non-drug targets. They applied a synthetic minority over-sampling technique to alleviate the class (target/non-target) unbalance problem. The authors used three different sets of features extracted from protein sequences, i.e. amino acid compositions, amino acid property group compositions and dipeptide composition, and achieved an accuracy of 85.3% using leave-one-out cross-validation. However, this approach looked at drug targets in a very general sense, without considering any specific disease associations.

In contrast,^10^ studied the predictive power of gene-disease association data for novel target identification. They benchmarked several models for their classification problem in a semi-supervised learning setting, and achieved an accuracy of 71% on the hold-out test set with an artificial neural network (ANN) model. The authors found that the key data types for therapeutic target prediction were the existence of an animal model, gene expression and genetic data, all coming from the Open Targets platform^11^. However, as the authors themselves point out, animal models with disease-relevant phenotypes are biased towards well-studied genes or diseases, limiting the range of potential targets that will be considered. Furthermore, by including any type of disease, opportunities for utilizing disease-specific data or for drawing disease-specific conclusions may be missed.

In this paper, we propose a computational pipeline that supports target identification in oncology. Within this context, our approach helps assess the specific value of different data types, as well as the benefits of combining multiple data types. Specifically, we investigate the computational prediction of novel therapeutic targets for anti-cancer therapy. We analyze the performance of five different machine learning classifiers: random forests (RF), artificial neural networks (ANN), support vector machines (SVM), logistic regression (LR), and gradient boosting machines (GBM). To train our models we gather and integrate gene mutation, expression and essentiality data, and complement our set of features with a numerical embedding of the interaction network of protein-coding genes. Using this approach we generate individual models for nine different cancer types, providing results that are disease-specific but cover a broad range of cancers. In each case we assess the contribution of each data type to aid model interpretation, and make predictions for more than 15,000 protein-coding genes. We thus produce genome-wide, unbiased ranked lists of putative novel targets per cancer type, available for follow-up experimental validation.

## Methods

### Overview

Figure 1 provides a graphical summary of the analytical approach implemented in this work. We applied the same method to each of the nine different cancer types that were considered. The analyses were performed in Python, primarily using the “Scikit-learn” package^12^. Fundamentally, we followed a supervised learning approach, with the goal of classifying human genes into “targets” and “non-targets” respective to a particular cancer type. By “targets” we mean genes that could reasonably be considered as therapeutic targets for drug development programs aimed at treating the corresponding cancer. We define a suitable dataset for model training and testing based on two sources of “known” (gold-standard) target genes: Targets of approved cancer drugs, and cancer driver genes. We reasoned that combining targets from both sources would help model generation by providing a larger and more comprehensive set of “positive” observations. Conceptually, the cancer-specific set of target genes is complemented by an equally-sized set of non-target genes, which are sampled at random from among the remaining human protein-coding genes. Having a dataset with balanced classes is generally advantageous for modeling and simplifies the interpretation of results, as both classes are equally likely *a priori*. In practice we pick ten different, non-overlapping sets of non-target genes, and average results from the ten resulting models, in order to reduce the variance of the classification.

**Figure 1.**
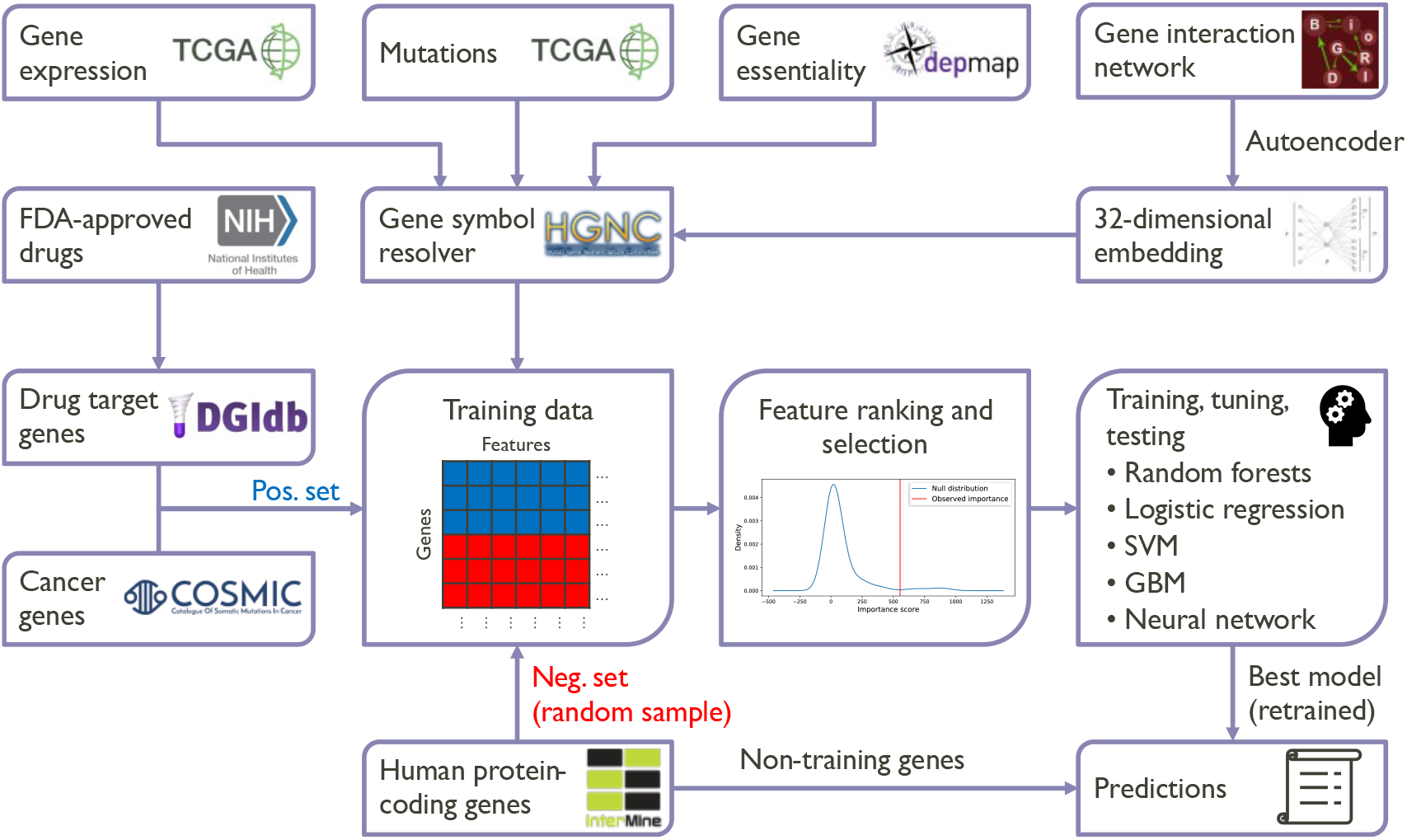
Graphical summary of the overall analytical approach followed in this work.

The target/non-target status of each gene in the dataset defines the class label that we want to predict. The features or attributes of the genes that models should learn to associate with the labels come from genome-wide datasets that are expected to be highly informative in this context - the same datasets that experts would interrogate to validate potential targets in oncology: From The Cancer Genome Atlas (TCGA), we use gene expression and mutation data, averaged over patient samples from the corresponding cancer cohort. From the Cancer Dependency Map (DepMap), we use gene essentiality data, in the form of average sensitivity scores from CRISPR knock-out experiments in cancer cell lines of the corresponding lineage. In separate analyses for each cancer type, these primary data types (expression, mutation, essentiality) are augmented by including gene-gene interaction data from BioGRID^13^ for human protein-coding genes.

The interaction network neighborhood of each gene was encoded in 32 numerical features using a neural network autoencoder^14^. We have recently shown that the information captured by such an embedding can be relevant for drug target identification^15^. All four data types combined give 35 features per gene and cancer type.

Before building machine learning models, we assessed whether any of these features showed a significant difference between the two classes by performing non-parametric significance tests. Each cancer type-specific dataset was then randomly split with a stratification strategy into a training set and a test set. For the following steps, only the training set data was used. In order to measure feature importance in a multivariate setting and select the best features for model generation, we applied a procedure based on the “random forests” machine learning algorithm. We then trained models on the selected features using five different machine learning methods: random forests, logistic regression, support vector machine, gradient boosting machine, and artificial neural network. Model hyperparameters were tuned by cross-validation to avoid overfitting.

We evaluated the performance of each model on the hold-out test set, and accumulated results over all ten random sets of non-targets. We chose the machine learning method that achieved the highest area under the ROC curve (AUROC) for our final model. We re-trained this model using the full (training + test) dataset. Again, this was done for each of the ten non-target sets, giving an ensemble of ten related models. Predictions for the target/non-target status of all human protein-coding genes were made using this ensemble, by averaging the predictions from the individual models. Finally, genes were ranked according to their predicted probability of being a potential target for cancer drug development.

### Dataset generation

#### Target and non-target genes

To generate a positive set of target genes for a specific cancer type, we first retrieved the list of FDA-approved drugs for this cancer type from the website of the US National Cancer Institute (https://www.cancer.gov/about-cancer/treatment/drugs/cancer-type). Next, we queried the Drug-Gene Interaction Database (DGIdb)^16^ for a list of target genes associated with each drug. To exclude untargeted chemotherapies and off-targets, only target genes with available “interaction type” information were included in the positive set. This set was extended by adding cancer driver genes implicated in the corresponding cancer type according to the Cancer Gene Census^17^. The nine cancer types considered in this study were selected based on the amount of available target data, as summarized in Table 1.

**Table 1.** Data availability summary for the data used in this work. Each column represents a different cancer type. In the first row the number of drugs is shown; second row depicts the target genes of these drugs; third row is the number of cancer genes from the Cancer Gene Census; fourth row shows the total number of “positive” genes and last row shows total number of “positive” genes with complete biological data across the cancer types in this study.

|  | Bladder | Breast | Colon | Kidney | Leukemia | Liver | Lung | Ovarian | Pancreatic |
| --- | --- | --- | --- | --- | --- | --- | --- | --- | --- |
| Drugs | 10 | 31 | 13 | 13 | 29 | 6 | 7 | 9 | 7 |
| Target genes | 26 | 58 | 32 | 31 | 99 | 26 | 11 | 31 | 41 |
| Cancer genes | 13 | 36 | 61 | 1 | 203 | 1 | 63 | 29 | 21 |
| Total genes | 39 | 94 | 93 | 32 | 302 | 27 | 74 | 60 | 62 |
| Genes with data | 39 | 87 | 83 | 32 | 228 | 27 | 67 | 57 | 55 |

In order to retrieve negative samples (non-target genes) for each cancer type, we queried HumanMine (https://www.humanmine.org, a database of human biological data based on the InterMine^18^ platform) for human protein-coding genes. From a pool of more than 17,000 genes, we sampled randomly without replacement to generate a negative set of equal size to the positive set. We made the reasonable assumption that true target genes will be very rare; thus, while we cannot guarantee that an (unknown) true target gene gets sampled by chance, such an occurrence should be highly unlikely. The process was repeated nine times to produce ten pairwise disjoint negative sets, in order to reduce the impact of each random sample on statistics and predictions.

#### Biomedical and genomic data

Preprocessed gene-level expression and somatic non-silent mutation data from The Cancer Genome Atlas (TCGA) pan-cancer cohort was downloaded via the UCSC Xena portal^19^ (dataset IDs: “*EB* + +*AdjustPANCAN_IlluminaHiSeq_RNASeqV*2” and “*mc*3.*v*0.2.8.*PUBLIC.nonsilentGene*”, version 2016-12-29). Gene essentiality data in the form of normalized sensitivity scores (CERES) from genome-scale CRISPR knock-out screens in cell lines was downloaded from DepMap^6^ (“*Achilles_gene_effect.csv*”, release 19Q2). Aggregation steps were necessary to transform this data to a per-cancer type format: Patient samples from TCGA were matched to cancer types using the available metadata. A per-cancer mutation rate was calculated for each gene as the mean of the data values (1/0 for mutated/non-mutated) across all corresponding samples. Similarly, gene expression per cancer type was calculated as the median expression value across corresponding samples. For the DepMap data, each cell line was matched to its cancer type of origin, and median sensitivity scores per cancer type and gene were calculated. Gene symbols were resolved using the REST API provided by the HUGO Gene Nomenclature Committee (genenames.org)^20^.

#### Gene-gene interaction network

We generated a representation of the gene-gene interaction network of all human protein-coding genes as follows: The list of protein-coding genes was exported from HumanMine^18^. For each gene, we queried the BioGRID database^13^ (version 3.5.171 from March 2019) for the genes interacting directly with it, and generated an edge list file for use with common computational frameworks. This resulted in a network comprising 17,389 nodes (corresponding to the protein-coding genes) and 323,247 edges. We then computed a 32-dimensional numerical embedding of the interaction network using sequence-based embedding with diffusion graphs^14^. Note that in contrast to the “primary” features, the resulting network embedding features were not specific to a cancer type.

#### Training and test sets

Each cancer type-specific dataset contained information on three “primary” features (mutation, expression, essentiality) and 32 network embedding features, pertaining to a number of “gold-standard” target genes and the same number of randomly chosen non-target genes. Each dataset was randomly split into a training set (70% of genes) and a test set (30% of genes), using stratified sampling to preserve the class balance (target/non-target) in both sets. Since ten separate sets of non-target genes were sampled for each cancer type, there were actually ten training and test sets for each case (including identical sets of target genes), so most analyses were repeated ten times and the results aggregated. The training sets were used for multivariate feature ranking and selection, and model training (including hyperparameter tuning). The test sets were used for assessing model performance and selecting the best model per cancer type. Both sets together were used for univariate feature rankings and retraining the selected models before making predictions.

### Univariate analysis

For each feature and cancer type, differences between feature values of target and non-target genes in the dataset were assessed using Mann-Whitney-Wilcoxon tests. In contrast to the multivariate feature selection methodology, the full dataset (training + test set) was used and all ten negative sets were pooled. Multiple testing correction was performed using the Benjamini-Hochberg procedure for controlling the false discovery rate.

### Feature selection methodology

We used a random forests (RF)-based permutation importance^21^ as our feature importance measure, which was calculated as follows for each training set: A null distribution for the importance of each feature was derived from the importance score of the RF model in a non-informative setting. To this end, we shuffled the labels (target/non-target) 100 times, trained a model and recorded the (information gain) importance scores in each shuffle. Each feature’s score in the “real” (non-shuffled) RF model was compared to the null distribution for that feature. The results were expressed as z-scores and averaged over the ten iterations on the negative sets. Only features with average z-scores of 0.5 or higher were retained for model training.

### Model generation

Based on the cancer type-specific training sets and the selected features, we trained models using five different machine learning methods. We performed 5-fold cross-validation to tune the following model hyperparameters. For random forests: the maximum tree depth, number of trees per forest, and maximum number of features to be considered at each split. For the support vector machine: the kernel function (linear or RBF), cost parameter, and kernel bandwidth (RBF kernel only). For the gradient boosting machine: the learning rate, as well as the three decision tree parameters (see random forests). The logistic regression model required no parameter tuning. We utilized the Keras library^22^ for the implementation of the neural network, with an architecture as described in Supplementary Table S1. The dropout mechanism (probability = 0.5) and batch normalization were used to regularize the model and speed up the training process, respectively. Processing was performed using GPU acceleration on a workstation with an NVIDIA GeForce GTX 1050 GPU and 16GB of RAM. The hyperparameter search space for each of the methods is shown in Supplementary Table S2. After models were evaluated on the test sets (see below), the best-performing machine learning method was selected for each cancer type; models were then retrained on the whole dataset (training + test) before making predictions.

### Model evaluation

Once the models are trained, we assessed their generalization performance on an independent test set (see subsection “Training and test sets” above), and selected the best performing method for each cancer type. In each case, models were evaluated by averaging their results on the test set across ten repeats of randomly sampling the “non-target” gene set. We used a typical classification metric to evaluate prediction performance - the area under the ROC curve (AUROC). This measure quantifies the probability that the classifier will rank a randomly chosen positive data point higher than a randomly chosen negative one. To assess the quality of predictions made for unlabeled genes, we considered the number of unique publications linking the gene and respective cancer type, retrieved from Open Targets^11^, as an orthogonal source of validation.

## Results

### Feature importance

In order to understand which data types were most informative for predicting new therapeutic targets in different cancers, we analyzed the importance of features in our data in a univariate (one feature at a time) and multivariate (all features used together) setting. In the univariate case, we used a statistical test to quantify whether feature values differed significantly between target and non-target genes. Figure 3 (A) summarizes the distribution of adjusted p-values across cancer types for each feature. We observe that gene expression, mutation, and some of the network embedding features were consistently significant (with few exceptions). However, the majority of the embedding features as well as gene essentiality varied widely in importance and were rarely significant. Comparing feature importance profiles across cancer types revealed generally low correlations (Figure 3, B).

For the multivariate case, we used a robust approach to analyze the contribution of each feature in context with the other features. Figure 2 shows two examples of feature importance relative to a corresponding null distribution, illustrating our approach for measuring multivariate feature importance in terms of z-scores.

**Figure 2.**
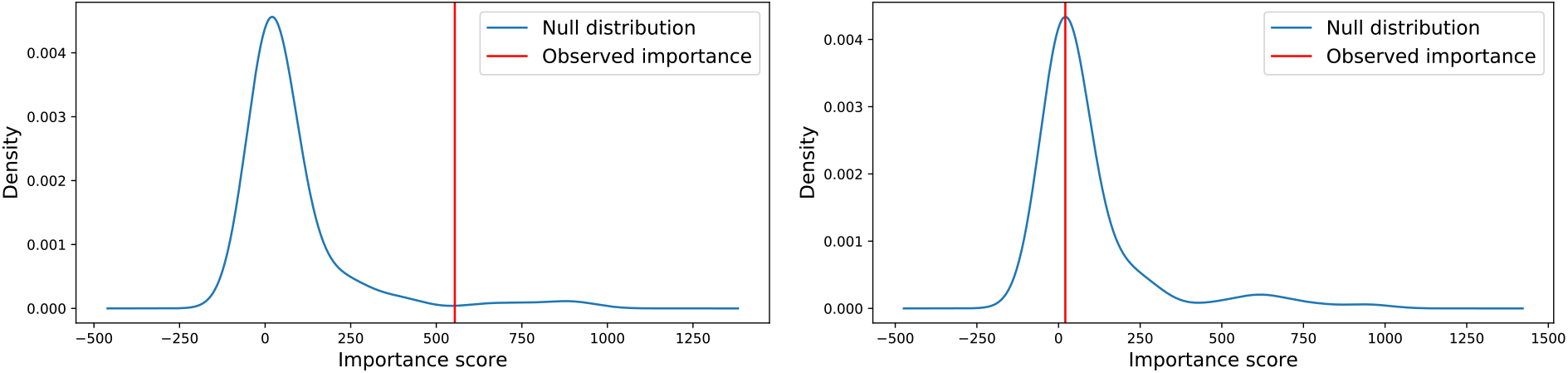
Illustrative examples of assessing multivariate feature importance by comparing to a null distribution. Data from lung cancer. Top: The “mutation” feature is shifted to the right of its null distribution, indicating high importance. Bottom: The “essentiality” feature appears close to the mean of its null distribution, making this a bad predictor.

These Z-scores are summarized in Figure 3 (C). Two network embedding features appear as most informative overall, followed by mutation, another embedding feature, and expression. There is high agreement between the univariate and multivariate rankings of the highly informative (top 6) features (Figure 3, A/C). Somewhat surprisingly, the gene essentiality feature was never ranked above our significance threshold (z-score of 0.5) in any cancer type. On the other side, it is very clear that certain of the network embedding features were consistently valuable for the target prediction task.

**Figure 3.**
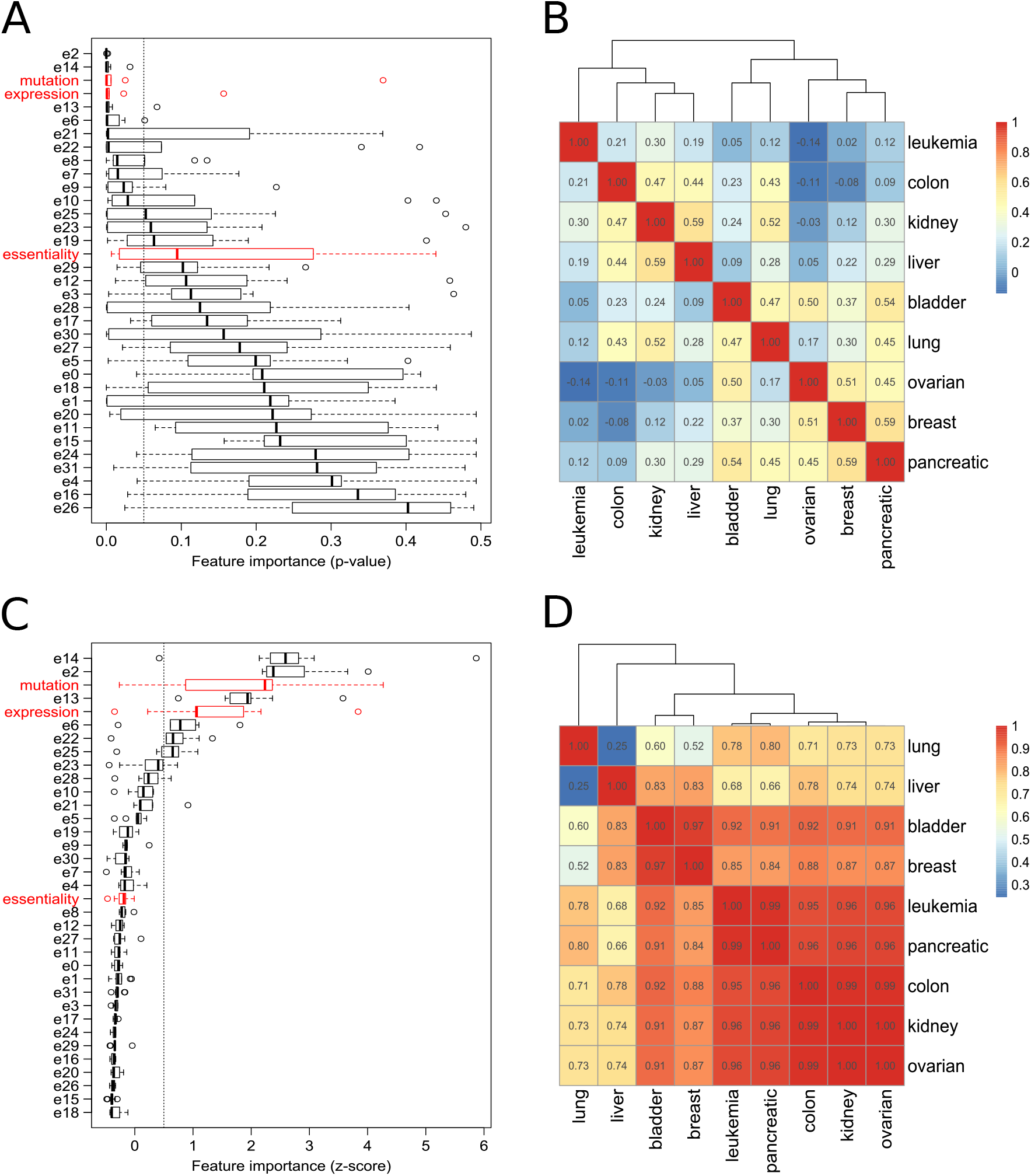
Univariate (A) and multivariate (C) feature rankings. Left column: Distribution of importance values for each feature across cancer types. “Primary” features are shown in red, network embedding features are in black. Lower p-values (A) and higher z-scores (C) mean higher importance. A p-value cut-off of 0.05 (A) and z-score cut-off of 0.5 (C) are indicated by dashed vertical lines. Right column: Correlation of feature importance values across the nine cancer types in the univariate (B) and multivariate (D) cases.

In contrast to the univariate case, we observe that the multivariate feature importance profiles are highly correlated between cancer types (Figure 3, D), with the exception of lung and liver cancer. Consistent with Figure 3 (C), in most cancer types a combination of network embedding features, mutation and expression was found to be important. However, in the liver cancer data only network embedding features scored highly; for lung cancer, mutation and expression (top 1 and 3 features) were more important than in other cancers. High mutation rates are typical for lung cancer, consistent with chronic mutagenic exposure caused by tobacco smoking^23^.

### Model performance

We initially trained machine learning models exclusively on the three “primary” cancer type-specific features: gene expression, mutation and essentiality. Model performance was evaluated on the test sets; the results for the best-performing models per cancer type are shown in Figure 4 (left).

**Figure 4.**
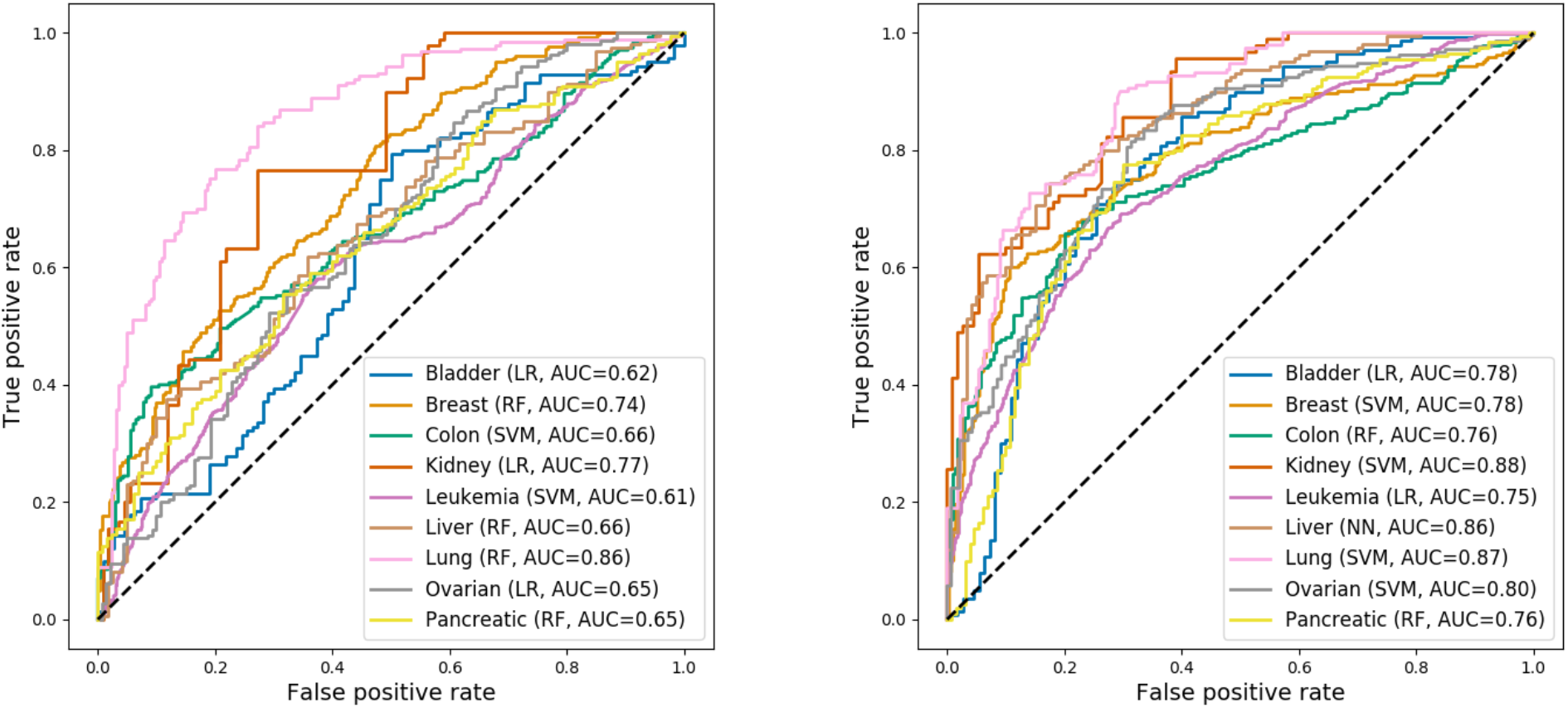
Generalization performances on the test sets for the best models across cancer types, measured in terms of AUROC. Left: Models trained on the three “primary” features (mutation, expression, essentiality). Right: Models trained after integrating the network embedding features and performing feature selection. The dashed lines correspond to a theoretical null model that predicts a label at random (AUROC=0.5).

The predictive performance achieved for different cancer types varied considerably. Despite the low number of features used for modeling, we observed respectable outcomes in three cancer types: lung, kidney and breast cancer, with AUROCs of 0.86, 0.77 and 0.74, respectively. In the remaining cancer types test set accuracies were low (AUROC 0.61-0.66), i.e. close to the baseline of 0.5 (random guess).

After including network embedding features in the datasets, performing feature selection, and training and evaluating new sets of models, we observed the performance metrics shown in Figure 4 (right). The predictive power of all models improved compared to the “primary” feature set, but the magnitude of improvement was variable. In general, previously low-performing models benefited more from the inclusion of additional features. The smallest improvement was observed for the lung cancer model. Notably, the “extended” models achieved test set AUROCs above 0.75 for all cancer types. AUROCs close to 0.9 were reached in three cases (kidney cancer: 0.88, lung cancer: 0.87, liver cancer: 0.86). The leukemia model achieved the lowest prediction performance (AUROC 0.75) on its test set, despite having by far the largest number of observations for training. We hypothesize that this is due to the considerable diversity of leukemias, which was not adequately represented in our dataset, where all subtypes were combined.

While the best-performing models for different cancer types came from different machine learning methods, the various methods generally achieved similar performance levels on the same dataset (Supplementary Table S3). As an exception, the gradient boosting machine performed less well in this study, presumably due to the relatively low number of observations available for training. Despite being simple linear classifiers, the logistic regression models were competitive with more complex models.

#### Prediction of novel therapeutic targets

We proceeded to carry out a genome-wide predictions using the best model for each cancer type, in order to identify novel targets that might be of therapeutic interest. Predictions were made for all human protein-coding genes with available data, except those that were part of the corresponding “positive” set of target genes used for model training and testing. For genes that were part of one of the ten “negative” (non-target) sets, only predictions from the other nine sets were considered. Predictions were made for almost 15,500 genes for each cancer type, except leukemia, where almost 13,600 predictions were made due to lack of TCGA expression data for some genes. The predicted target probabilities for different cancer types followed different distributions, but these could be aligned by transforming the predicted probabilities from each model into z-scores (see Supplementary Figure S1).

While we would expect only a minuscule fraction of genes to be valid therapeutic targets for any cancer type, due to the use of balanced training sets (with equal numbers of targets and non-targets), our models classified large fractions of genes as potential targets when a conventional probability cut-off of 0.5 was used (see Supplementary Table S4). However, our interest is in the ranking of genes, so calibrating the probabilities was not a concern.

We focus our discussion on the top five predictions for three of the cancer types - those with the highest numbers of known targets available for training (see Table 1). Tables 2, 3 and 4 show the five genes with highest predicted probabilities of being a target for leukemia, colon cancer and breast cancer, respectively. Top five predictions for the remaining cancer types are included as Supplementary Tables S5-S10.

**Table 2.** Top 5 predictions for leukemia.

| Gene | Full name | Probability | Citations |
| --- | --- | --- | --- |
| SNHG29 | Small Nucleolar RNA Host Gene 29 | 0.979 | N/A |
| HNRNPU | Heterogeneous Nuclear Ribonucleoprotein U | 0.961 | 3 |
| RPS6KA2 | Ribosomal Protein S6 Kinase A2 | 0.935 | 2 |
| LCOR | Ligand Dependent Nuclear Receptor Corepressor | 0.915 | 2 |
| ARHGAP27 | Rho GTPase Activating Protein 27 | 0.903 | 1 |

As a source of orthogonal validation, we obtained the number of articles in the scientific literature that linked our top predictions to the corresponding disease by querying Open Targets^11^. To check whether there was an overall association between predicted probability of being a target and the number of supporting citations, we randomly sampled 1000 genes per cancer type and compared their target probabilities and citation counts. We found moderate, but statistically significant (p < 0.01 after Benjamini-Hochberg correction) Spearman correlations between both measures for half of the cancer types (bladder: 0.10, breast: 0.15, lung: 0.13, ovarian: 0.16, pancreatic: 0.13).

Our list of top predictions for leukemia targets is shown in Table 2. There was no disease information at all available in Open Targets for the top gene, SNHG29. However, under the alias LRRC75A-AS1 this gene is part of a three-lncRNA signature that has been identified for the prognosis prediction of acute myeloid leukemia (AML) patients^24^. The four other genes had some reported association with leukemia. For example, the RSK gene family, to which RPS6KA2 belongs, was identified to play an important role in maintaining AML cell survival and proliferation, positioning it as a promising target for AML treatment^25^. Furthermore, in a recent study^26^, LCOR was implicated in the progression of B-cell acute lymphoblastic leukemia. The evidence for ARHGAP27 is weak (mRNA expression in chronic lymphocytic leukemia), but the ARHGAP gene family has been linked to carcinogenesis through the dysregulation of Rho/Rac/Cdc42-like GTPases^27^.

Among the predictions for colon cancer targets (Table 3), the top gene, LRRK1, was not supported by text mining data from Open Targets. However, the mouse ortholog of this gene has been identified as a potential colorectal cancer (CRC) driver in a mutagenesis screen, and LRRK1 is frequently deleted in human CRC^28^. Surprisingly, even though MET is a known colon cancer oncogene^29^ with at least 156 publications relating it to the pathology, it was not marked as a target gene in either of the data sources we used to obtain the positive set of genes (at the time of analysis). However, it was picked up by our model as a top target for colon cancer. The next gene in the list, STAT5B, was found to have a statistically significant association with colon cancer risk^30^. In another study, RASA1 expression was shown to be regulated by the oncogene miR-21, promoting invasion and tumor formation ability in colon cancer RKO cells^31^. Finally, the INSR gene has prognostic relevance for CRC^32^; importantly, it is also being targeted by a clinical trial in phase 4 for that disease (ClinicalTrials.gov, ID NCT02032953).

**Table 3.** Top 5 predictions for colon cancer.

| Gene | Full name | Probability | Citations |
| --- | --- | --- | --- |
| LRRK1 | Leucine Rich Repeat Kinase 1 | 0.936 | 0 |
| MET | MET Proto-Oncogene, Receptor Tyrosine Kinase | 0.935 | 156 |
| STAT5B | Signal Transducer and Activator of Transcription 5B | 0.933 | 2 |
| RASA1 | RAS P21 Protein Activator 1 | 0.930 | 7 |
| INSR | Insulin Receptor | 0.924 | 9 |

The five most probable targets for breast cancer as predicted by our models are listed in Table 4. There is some support for these predictions in the literature. The microRNA miR-10b has been reported^33^ to promote breast cancer cell proliferation, migration, and invasion through inhibition of the expression of the transcription factor TBX5 - our top prediction. Another study^34^ discovered that overexpression of ZFP57 could inhibit the proliferation of breast cancer cells by inhibiting the Wnt/β- catenin pathway. Expression levels of GATA4 have been linked to breast cancer progression^35^. INSM1 was found to be a novel and sensitive marker of tumors with neuroepithelial differentiation, such as breast cancer^36^.

**Table 4.** Top 5 predictions for breast cancer.

| Gene | Full name | Probability | Citations |
| --- | --- | --- | --- |
| TBX5 | T-Box Transcription Factor 5 | 0.967 | 2 |
| ZFP57 | ZFP57 Zinc Finger Protein | 0.941 | 1 |
| GATA4 | GATA Binding Protein 4 | 0.934 | 1 |
| NEUROD6 | Class A Basic Helix-Loop-Helix Protein 2 | 0.931 | 0 |
| INSM1 | INSM Transcriptional Repressor 1 | 0.921 | 1 |

Overall, these results demonstrate that some of our top predictions are well supported by orthogonal evidence from text mining data. This suggests that our prediction framework could be beneficial for new therapeutic target identification in oncology.

## Discussion

In this work we have analyzed how computational intelligence methods can be applied to predict novel therapeutic targets in oncology. We compared five machine learning classifiers for the task of drug target prediction in nine different cancer types. Very simple models, incorporating only gene expression, mutation and essentiality information for individual genes, already achieved reasonable prediction performance for some cancer types (up to 0.86 AUROC for lung cancer). However, through the integration of gene-gene interaction data via network embedding features, combined with a robust feature selection approach, well-performing models could be generated for all nine cancer types (AUROCs between 0.75 for leukemia and 0.88 for kidney cancer). A caveat associated with the inclusion of interaction network data is the bias towards high connectivity of well-studied genes. We have to assume that the drug target and cancer driver genes at the core of this study fall into this group, hence we cannot rule out a corresponding bias in our models.

Another possible source of bias is the overrepresentation of certain enzyme families (especially kinases) among the targets of currently approved anti-cancer drugs. While our models do not directly incorporate information on protein function, they can only learn patterns that are represented in the training data, which may limit their ability to accurately predict targets from novel classes (e.g. RNA-modifying enzymes). Note that potential biases in a training dataset will affect any computational approach based on such data.

Our analysis of multivariate feature importance showed that a combination of mutation, gene expression and certain of the network embedding features was informative for target prediction in most cancer types. Unfortunately an interpretation of the important network features in terms of biological or graph-based properties (e.g. connectivity, clustering) is currently not possible, due to the neural network approach used to calculate the embedding. Surprisingly, DepMap-based gene essentiality was not estimated to be an important feature, although the underlying experimental data provides direct functional readouts for the effect of losing specific genes on proliferation in cancer cells. It is possible that our encoding of this information (as the average of the sensitivity scores) was not optimal for our purpose, and that a different encoding (e.g. the fraction of cell lines that are sensitive based on a predefined score cutoff) would have been more informative.

We applied our models to more than 15,000 protein-coding genes, in each case predicting the probability of the gene being a potential cancer type-specific target. In three example cancer types, we provided external evidence from the scientific literature, linking our top five predictions to the specific pathology. However, the level of literature support for our predictions varied, and generally the correlations between predicted probabilities and the number of supporting citations were low.

Constructing a validation procedure for predictions of drug target candidates is inherently problematic. Essentially, validating a proposed target would require the development and testing of a corresponding drug, which is not feasible at scale. Text mining of the scientific literature or of clinical trial information can provide orthogonal validation, but both depend on what has previously been studied. Simpler effects, such as response of a cell line to gene knock-out, are more easily validated experimentally, but it is questionable how well such results translate to human patients. The lack of importance of the gene essentiality feature in our analyses can be seen as an indicator of this uncertainty.

Despite some limitations, our study proves that computational intelligence methods can be used to discriminate known cancer type-specific target genes from non-target genes with good accuracy. It would be straightforward to extend our approach by integrating additional data types, such as copy number variation, DNA methylation or protein functional/structural information, in the future. As mentioned above for gene essentiality, more elaborate encodings for the current data types could be envisioned as well, in order to capture additional aspects besides an “average per gene”. Furthermore, while we only tested supervised learning algorithms here, semi-supervised learning methods could be useful alternatives. A semi-supervised approach would require a careful distinction between non-target genes and genes of unknown status, but could draw on additional information from the latter set. Future extensions notwithstanding, our work provides a novel way to generate hypotheses for therapeutic targets in oncology.

## Acknowledgements

AB acknowledges funding from Innovate UK (grant KTP011266). The authors would like to thank Julie Sullivan for InterMine support, Yaara Ofir-Rosenfeld for helpful discussions on target validation in oncology, and Gos Micklem and Yo Yehudi for valuable feedback on the manuscript.

## Author contributions statement

AB designed and implemented the software, performed the analyses and co-wrote the manuscript. DL supervised the project and contributed to the manuscript. HW conceived and supervised the project and co-wrote the manuscript. All authors read and agreed to the final manuscript.

