## Supplementary Material for "Genome-wide investigation of gene-cancer associations for the prediction of novel therapeutic targets in oncology"

by

Adrián Bazaga, Dan Leggate and Hendrik Weisser

Supplementary Table S1: Details of the artificial neural network architecture used in this work.

| Layer | Type | Number of neurons (output) | Activation function |
|---|---|---|---|
| 1 | Fully-connected | 64 | ReLU |
| 2 | Fully-connected | 128 | ReLU |
| 3 | Fully-connected | 16 | ReLU |
| 4 | Fully-connected | 2 | Sigmoid |

Supplementary Table S2: Hyperparameter search space for each of the machine learning methods

| Method | Parameters and range of values |
|---|---|
| Random Forest | Max depth = [4, 5, 6, 7, 8]<br># estimators = [300, 500, 700, 850, 1000]<br>Max features = [sqrt(# features), log2(# features), 30%, 50%] |
| Support Vector Machine | Kernel function = [linear, rbf].<br>For RBF kernel, gamma = [1e-3, 1e-4], C = [1, 10, 100, 1000].<br>For linear kernel, C = [1, 10, 100, 1000] |
| Gradient Boosting Machine | Learning rate = [0.005, 0.1].<br>Max depth = [4, 5, 6, 7, 8].<br># estimators = [300, 500, 700, 850, 1000].<br>Max features = [sqrt(# features), log2(# features), 30%, 50%] |
| Logistic Regression | N/A |

Supplementary Table S3: Performance in terms of test set AUC achieved by each of the five different machine learning methods across cancer types.

| Cancer type / Method | Bladder | Breast | Colon | Kidney | Leukemia | Liver | Lung | Ovarian | Pancreatic |
|---|---|---|---|---|---|---|---|---|---|
| Logistic Regression | 0.78 | 0.77 | 0.69 | 0.86 | 0.75 | 0.84 | 0.81 | 0.79 | 0.75 |
| Support Vector Machine | 0.77 | 0.78 | 0.72 | 0.88 | 0.72 | 0.84 | 0.87 | 0.8 | 0.73 |
| Gradient Boosting Machine | 0.75 | 0.7 | 0.74 | 0.75 | 0.71 | 0.81 | 0.73 | 0.74 | 0.73 |
| Neural Network | 0.67 | 0.72 | 0.71 | 0.71 | 0.7 | 0.86 | 0.75 | 0.75 | 0.72 |
| Random Forests | 0.76 | 0.75 | 0.76 | 0.79 | 0.74 | 0.85 | 0.83 | 0.77 | 0.76 |

Supplementary Table S4: Total number of genes predicted as targets (probability $\geq 0.5$) by the best model for each of the cancer types

| Cancer type | Number of predicted targets |
|---|---|
| Bladder | 4473/15500 (28%) |
| Breast | 4129/15500 (26%) |
| Colon | 3246/15500 (20%) |
| Kidney | 5451/15500 (35%) |
| Leukemia | 4272/13600 (31%) |
| Liver | 3188/15500 (20%) |
| Lung | 4502/15500 (29%) |
| Ovarian | 4681/15500 (30%) |
| Pancreatic | 3750/15500 (24%) |

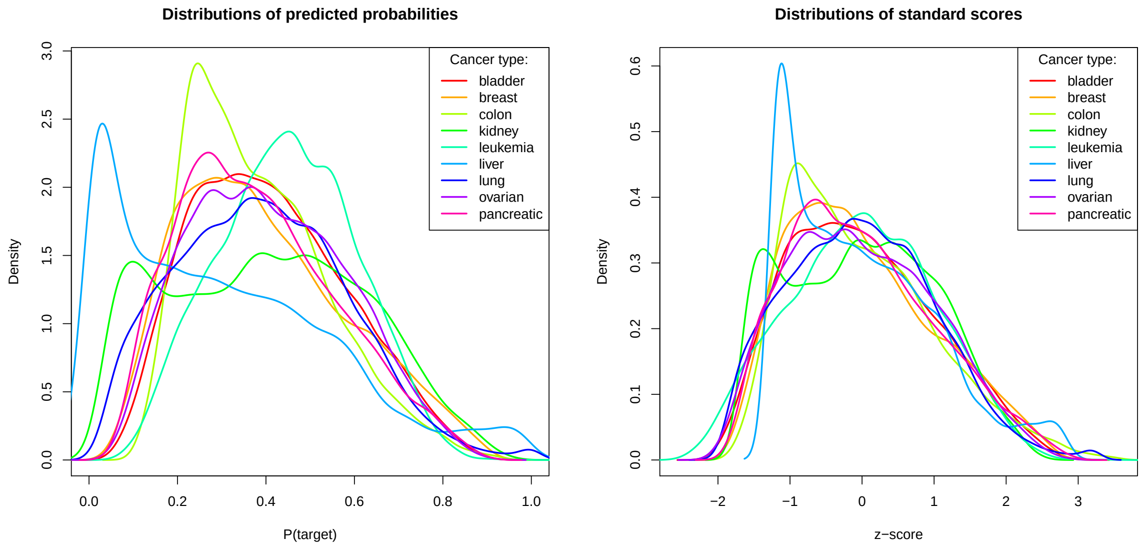


Supplementary Figure S1: Distributions (kernel density estimates) of genome-wide predicted probabilities for different cancer types, before (left) and after (right) scaling per cancer type.

Supplementary Table S5: Top 5 predictions for ovarian cancer.

| Gene | Full name | Probability | Citations |
|---|---|---|---|
| SLA | Src Like Adaptor | 0.919 | 0 |
| LY6E | Lymphocyte Antigen 6 Family Member E | 0.914 | 0 |
| TYROBP | TYRO Protein Tyrosine Kinase Binding Protein | 0.909 | 0 |
| JAK1 | Janus Kinase 1 | 0.908 | 0 |
| CCDC74A | Coiled-Coil Domain Containing 74A | 0.905 | 0 |

Supplementary Table S6: Top 5 predictions for pancreatic cancer.

| Gene | Full name | Probability | Citations |
|---|---|---|---|
| STAT1 | Signal Transducer And Activator Of Transcription 1 | 0.909 | 2 |
| PTPN12 | Protein Tyrosine Phosphatase Non-Receptor Type 12 | 0.901 | 1 |
| MYO1D | Myogenic Differentiation 1 | 0.899 | 1 |
| NBEAL2 | Neurobeachin Like 2 | 0.896 | 0 |
| INTS3 | Integrator Complex Subunit 3 | 0.895 | 1 |

Supplementary Table S7: Top 5 predictions for kidney cancer.

| Gene | Full name | Probability | Citations |
|---|---|---|---|
| TYROBP | TYRO Protein Tyrosine Kinase Binding Protein | 0.961 | 0 |
| SLA | Src Like Adaptor | 0.958 | 0 |
| PEAR1 | Platelet Endothelial Aggregation Receptor 1 | 0.956 | 0 |
| JAK1 | Janus Kinase 1 | 0.952 | 0 |
| KL | Klotho | 0.950 | 0 |

Supplementary Table S8: Top 5 predictions for bladder cancer.

| Gene | Full name | Probability | Citations |
|---|---|---|---|
| MDFIC | MyoD Family Inhibitor Domain Containing | 0.930 | 0 |
| PRDM2 | PR/SET Domain 2 | 0.908 | 0 |
| POU3F1 | POU Class 3 Homeobox 1 | 0.904 | 3 |
| HMGA1 | High Mobility Group AT-Hook 1 | 0.888 | 0 |
| PRSS8 | Serine Protease 8 | 0.887 | 2 |

Supplementary Table S9: Top 5 predictions for liver cancer.

| Gene | Full name | Probability | Citations |
|---|---|---|---|
| PEAR1 | Platelet Endothelial Aggregation Receptor 1 | 0.999 | 0 |
| SLA | Src Like Adaptor | 0.999 | 0 |
| SIT1 | Signaling Threshold Regulating Transmembrane Adaptor 1 | 0.998 | 2 |
| ZAP70 | Zeta Chain Of T Cell Receptor Associated Protein Kinase 70 | 0.998 | 0 |
| FCRL3 | Fc Receptor Like 3 | 0.998 | 0 |

Supplementary Table S10: Top 5 predictions for lung cancer.

| Gene | Full name | Probability | Citations |
|---|---|---|---|
| TENM1 | Teneurin Transmembrane Protein 1 | 1.0 | 0 |
| CCDC7 | Coiled-Coil Domain Containing 7 | 1.0 | 0 |
| NAA38 | N(Alpha)-Acetyltransferase 38, NatC Auxiliary Subunit | 1.0 | 0 |
| B3GNT2 | UDP-GlcNAc:BetaGal Beta-1,3-N-Acetylglucosaminyltransferase 2 | 1.0 | 0 |
| RPS6KA2 | Ribosomal Protein S6 Kinase A2 | 1.0 | 1 |
